# Thyroid hormones deficiency impairs male germ cell development: a cross talk between hypothalamic-pituitary-thyroid, and - gonadal axes in zebrafish

**DOI:** 10.1101/2022.01.30.478377

**Authors:** Maira S Rodrigues, Aldo Tovo-Neto, Ivana F da Rosa, Lucas B Doretto, Hamideh P Fallah, Hamid R Habibi, Rafael H Nóbrega

**Affiliations:** Aquaculture Program (CAUNESP), São Paulo State University (UNESP), 14884-900, Jaboticabal, São Paulo, Brazil; Reproductive and Molecular Biology Group, Department of Structural and Functional Biology, Institute of Biosciences, São Paulo State University (UNESP), 18618-970, Botucatu, São Paulo, Brazil; Department of Biological Sciences, University of Calgary, 500 University Drive NW, Calgary, Alberta, T2N 1N4, Canada

**Keywords:** hypothyroidism, methimazole, thyroid hormones, spermatogenesis, zebrafish

## Abstract

In vertebrates, thyroid hormones, including thyroxine (T4) and triiodothyronine (T3), are critical players in controlling different physiological processes such as development, growth, metabolism among others. There is evidence in mammals that thyroid hormones are also an important component of the hormonal system that controls reproduction, although studies in fish remain poorly investigated. Here we tested this hypothesis by investigating the effects of methimazole-induced hypothyroidism on the testicular function in adult *D. rerio*. Treatment of fish with methimazole, *in vivo*, significantly affected the progression of zebrafish spermatogenesis by inducing the accumulation of pre-meiotic cells, delaying cell differentiation and meiosis, as well as reducing the number of spermatozoa. The observed impairment of spermatogenesis by methimazole was correlated with significant changes in transcript levels for several genes involved in the control of reproduction. Using an *in vitro* approach, we also demonstrated that in addition to affecting the components of the brain-pituitary-peripheral axis, T3 also exerts direct action at the level of the testis. These results support the hypothesis that thyroid hormones are an essential component of multifactorial control of reproduction and testicular function in zebrafish and possibly other vertebrates.

## 1 Introduction

The production of thyroid hormones in fish and other vertebrates is under the control of the hypothalamic-pituitary–thyroid (HPT) axis (Cooke et al., 1994; Tousson et al., 2011; Duarte-Guterman et al., 2014; Kang et al., 2020). The thyrotropin (TSH)-releasing factor [thyrotropin-releasing hormone (TRH)/corticotropin-releasing hormone (CRH)] stimulates the anterior pituitary to release TSH, which in turn, promotes the synthesis and release of thyroid hormones, including T4 and T3, by thyroid follicles (Larsen et al., 1998). Of the two, T3 is the more biologically active thyroid hormone owing to its affinity for the nuclear thyroid hormone receptor (Carr and Patiño, 2011). The HPT axis acts parallel to the hypothalamic-pituitary–gonadal (HPG) axis, which involves a large number of hormones, including the gonadotropin-releasing hormone (GnRH) that stimulates the secretion of gonadotropin hormones, luteinizing hormone (LH), and follicle-stimulating hormone (FSH) (Schulz et al., 2010; Pankhurst, 2016) which are crucial for testis development and spermatogenesis in fish (Schulz et al., 2010; for a review see Xie et al., 2020). There is evidence for interaction between HPT and HPG axes in vertebrates (Teerds et al., 1998; Ariyaratne et al., 2000; Matta et al., 2002; Wagner et al., 2009; Morais et al., 2013; Kang et al., 2020), although this subject remains poorly investigated in fish, particularly, in the male reproductive system (Habibi et al., 2012; Castañeda-Cortés et al., 2014; Tovo-Neto et al., 2018; Ma et al., 2020a, b). In mammals, it has been shown that T3 regulates the growth and maturation of testis by inhibiting immature Sertoli cell proliferation and stimulating their terminal differentiation (Cooke and Meisami, 1991; Hess et al., 1993; Cooke et al., 1994). Furthermore, in postnatal rat testis, an important action of thyroid hormones is to initiate the onset of Leydig cell differentiation and stimulation of steroidogenesis (Ariyaratne et al., 2000), in part, by increasing the expression of steroidogenic acute regulatory protein (StAR) (Mendis-Handagama and Ariyaratne, 2005). The presence of thyroid hormone receptors in the mammalian testis, particularly in Leydig cells, suggests both direct and indirect actions of thyroid hormones on testicular function (Hernandez, 2018). The regulatory role of thyroid hormones is complex, species specific, and dependent on developmental stages. Neonatal hypothyroidism was shown to impair testicular growth and sperm production in rats (França et al., 1995), hamsters (*Mesocricetus auratus*) (Jansen et al., 2007), and juvenile teleost fish (*Oreochromis niloticus*) (Matta et al., 2002).

A number of researchers have investigated the role of thyroid hormones in fish embryogenesis, larval development, and growth (Blanton and Specker, 2007; Mukhi et al., 2007; Orozco et al., 2012). With regards to fish male reproduction, there are some evidence that thyroid hormones can affect spermatogenesis (Cyr and Eales, 1996; Wagner et al., 2009; Nelson et al., 2010; Habibi et al., 2012; Morais et al., 2013; Safian et al., 2016; Tovo-Neto et al, 2018). In adult catfish, *Clarias gariepinus*, treatment with thiourea (a thyroid disruptor) reduced 11-ketostestosterone (11-KT) and testosterone (T) production (Swapna et al., 2006), leading to male reproductive system disruption. Morais and collaborators (2013) demonstrated the influence of thyroid hormones on zebrafish spermatogenesis using an *ex vivo* approach. In the same study, the authors showed that T3 stimulated the increase in mitotic index of type A undifferentiated spermatogonia (A_und_) and Sertoli cells through Igf3 (Insulin-like growth factor like 3), a Sertoli cell stimulatory growth factor (Morais et al., 2013). Moreover, the same authors showed that T3 potentiated FSH actions on steroid release and enhanced Fsh-stimulated *cyp17a1* (17α-hydroxylase/17,20 lyase) and *ar* mRNA levels in zebrafish testis (Morais et al., 2013). There is also evidence that thyroid hormones interact with other reproductive peptides such as GnRH and gonadotropin-inhibitory hormones (GnIH) *in vivo* (Ma et al., 2020a, b) and *in vitro* in the zebrafish testis (Rodrigues et al., 2021). Other studies have demonstrated that GnRH stimulates thyroid activity in a freshwater murrel, *Channa gachua*, and two carp species, *Catla catla* and *Cirrhinus mrigala* (Roy et al., 2000). GnRH injection increases plasma T4 levels in different vertebrate species (Jacobs et al., 1988; Roy et al., 2000; Chiba et al., 2004), suggesting an effect of endogenous pituitary gonadotropin release due the heterothyrotropic activities of GnRH on the thyroid (Mackenzie, 1982, 1987). In general, these findings support the hypothesis that normal thyroid hormone action is critical for HPG axis function and normal gonadal function. However, significant gaps remain regarding exact physiological significance of thyroid hormones on fish male reproductive function.

The aim of this study was to further investigate the influence of thyroid hormones on male zebrafish (D. *rerio*) reproduction using *in vivo* and *ex vivo* approaches. We first evaluated the effect of hypothyroidism induced by methimazole and co-treatment with T4 on zebrafish spermatogenesis by histomorphometrical measurement and measured testicular transcript levels for genes related to reproduction, as well as 11-KT plasma levels and basal and FSH-induced 11-KT release *in vitro*. Subsequently, we investigated the effects of hypothyroidism induced by methimazole treatment and T3 injection, on zebrafish brain and pituitary by transcript measurement. Finally, we investigated long-term effects of T3 on zebrafish spermatogenesis by histomorphometrical analysis, and transcript levels of a selected genes. The results provide a framework for better understanding of the role of thyroid hormone in the control of male reproduction in zebrafish.

## 2 Methods

### 2.1 Animals

Sexually mature male zebrafish (4–5-months-old) were bred and raised in the aquarium facility of the Department of Structural and Functional Biology, Institute of Biosciences, Botucatu, São Paulo State University (UNESP). The animals were kept in 6-L tanks in the recirculating system under constant temperature conditions, similar to the natural environment (28 °C), and proper photothermal conditions (14-h light/10-h dark). pH, salinity, dissolved oxygen, and ammonia were monitored in all tanks every other day. Fish were fed twice a day with commercial food (Sera Vipan Flakes®). Handling and experimentation were performed according to the Brazilian legislation regulated by National Council for the Control of Animal Experimental (CONCEA) and Ethical Principles in Animal Research (Protocol n. 1031-CEUA) and University of Calgary animal care committee and in accordance with the guidelines of the Canadian Council of Animal Care (Protocol #AC19-0161).

### 2.2 Treatment solutions: methimazole-induced hypothyroidism and reversal treatment with T4

A solution of methimazole (1-methyl-3H-imidazole-2-thione) (CAS 60-56-0; MW, 114.17 g/mol; purity, ≥99%; Sigma-Aldrich) was used to induce hypothyroidism. A stock solution of methimazole was prepared by dissolving 11.4 g of methimazole in 1000 mL of distilled water. Exposure concentration of 1 mM was prepared by mixing 10 mL of the stock solution per liter of water in the tank. A stock solution (1 mg/mL) of T4 (CAS 51-48-9; MW, 776.87 g/mol; Sigma-Aldrich) was prepared by dissolving 1 mg of T4 in 1 mL of DMSO (dimethyl sulfoxide; CAS 67-68-5; MW, 78.13 g/mol; Vetec), and a treatment solution of 100 μg/L was prepared by adding 100 μL of the stock solution per liter of reconstituted water. To avoid problems while adding a new variable, the same amount of DMSO was added in all the tanks. Two thirds of the water volume was replaced every day (i.e., semi-static aquatic exposure). The working concentration of methimazole (1 mM) and T4 (100 μg/L) were selected based on previous studies (Sharma and Patiño, 2013; Sharma et al., 2016; Rodrigues et al., 2021). In this study, adult male zebrafish (n = 144) were divided into four replicate tanks per experimental group: control [only filtered water (n = 36)]; group I [filtered water with T4 (100 μg/L) (n = 36)]; group II [filtered water with methimazole (1 mM) (n = 36)]; group III [1 mM of methimazole followed by addition of T4 (100 μg/L) (n = 36) as reversal treatment group (methimazole + T4)]. In the T4 group (group I), males were exposed to reconstituted water containing 100 μg/L of T4 from the second week until the end of treatment. In the methimazole group (group II), adult males were exposed to reconstituted water containing 1 mM methimazole for 3 weeks. Also, the reversal treatment (methimazole + T4) group (group III), males were exposed to reconstituted water containing 1 mM methimazole for 3 weeks, and T4 (100 μg/L) was added in the water from the second week until the end of treatment. The reversal treatment with T4 was performed to address whether the effects observed were a direct effect of the lowering/removal of thyroid hormones. The experimental groups were euthanized by overdose with benzocaine hydrochloride (250 mg) previously dissolved in ethanol and then mixed with water. After euthanasia, the heads (control, methimazole and methimazole + T4) were collected for histological analysis of the thyroid follicles, while the testes from control, methimazole, and methimazole + T4 groups were dissected and immediately used for *in vivo* experiments (histomorphometric analysis and gene expression); androgen plasma levels and *ex vivo* organ culture experiment [short-term (18 h) incubation for androgen release by zebrafish testicular explants] were available on the treatments (control, T4, methimazole and methimazole + T4 groups) (see below). The brain and pituitary from the control and methimazole groups were collected for gene expression (see below).

### 2.3 Thyroid follicles histology

Thyroid follicles from control, methimazole and methimazole + T4 groups were examined histologically. Thyroid follicles appear dispersed among the afferent branchial arterioles (Patinõ et a., 2003; Van der Ven et al., 2006). Head was separated from the trunk, dissected, and fixed in modified Karnovsky (2% glutaraldehyde and 4% paraformaldehyde in Sorensen buffer [0.1 M, pH 7.2]) for at least 24 h at room temperature. After, samples were dehydrated, embedded in Technovit (7100) historesin (Heraeus Kulzer, Wehrheim, Germany), and serial sections at 3-μm thickness were obtained and stained with 0.1% toluidine blue in 1% sodium borate, according to conventional histological procedures. Then, histological sections were examined and documented using a Leica DMI6000 microscope.

### 2.4 Histomorphometrical evaluation of zebrafish spermatogenesis

After adult zebrafish were exposed to methimazole and methimazole + T4, testes (n = 8 per treatment) were dissected and immediately fixed in 4% Karnovsky fixative at 4°C overnight, dehydrated, embedded in Technovit (7100) historesin (Heraeus Kulzer, Wehrheim, Germany), sectioned at 3-μm thickness, and stained with 0.1% toluidine blue, according to conventional histological procedures. The slides were evaluated, and the proportion of section surface area of spermatogenic cysts containing type A undifferentiated spermatogonia (A_und*_ and A_und_), type A differentiated spermatogonia (A_diff_ type B spermatogonia (SpgB), spermatocytes (Spc), and spermatids (Spt) were determined. Intersection points were counted on the histologic fields, for which five fields per slide (n = 8 slides per treatment) were quantified using a grid of 540 (54 × 10) intersections under 100x objective lens. The proportion of section area occupied by different germ cell types were represented as fold-change of control value.

The relative number of spermatozoa per area was quantified for each treatment. Twenty different fields were captured from a 100x objective lens and analyzed by IMAGEJ Software (available at http://imagej.nih.gov/ij/index.html) for quantification of spermatozoa number according to Fallah et al., (2019, 2020), Tovo-Neto et al., (2020) and Rodrigues et al., (2021). This software allows quantification of several types of cells by limiting the size of the particle. First, it is necessary to defocus the image, and this filter may avoid false positive cells and improve the measurement. Next, threshold feature is activated, which separates the background comprising unspecific particles of interest. Then, the RGB image is converted into gray scale picture (8 or 16 bytes), and the threshold allows for lower intensity signals to be white and higher intensity signals to be black. The algorithm (for analyzing particles) can be applied for measuring black particles in the range of pre-fixed size.

### 2.5 Transcript analysis by quantitative real-time PCR (qPCR)

Total RNA from testes (control, methimazole, and methimazole + T4 groups) was extracted using TRIzol™ (Invitrogen, Carlsbad, CA, USA), according to the manufacturer’s instructions, and quantity and purity were checked with a NanoDrop™ One Spectrophotometer (Thermo Scientific, Madison, WI, USA). cDNA synthesis was performed as described previously (Nóbrega et al., 2010). qPCR reactions were conducted using 5 μL of 2x SYBR-Green Universal Master Mix, 1 μL of forward primer (9 mM), 1 μL of reverse primer (9 mM), 0.5 μL of DEPC water, and 2.5 μL of cDNA. The relative mRNA levels of *thrα* and *thrβ* (thyroid hormone receptors), *fshr* (follicle-stimulating hormone receptor), *cyp17a1* (17α-hydroxylase/17,20 lyase/17,20 desmolase), *insl3* (insulin-like peptide 3), *cx43* (testicular connexin), *igf3* (insulin-like growth factor 3), *amh* (anti-Müllerian hormone), *gsdf* (gonadal somatic cell derived factor), *nanos2* (marker of undifferentiated spermatogonia), *dazl* (deleted in azoospermia-like), *sycp3l* (synaptonemal complex protein 3) and *odf3a* (outer dense fiber protein 3) were measured in the different treatments.

Brain (n = 8) and pituitary (n = 4 pools of 4 pituitaries for each pool) were collected from control and methimazole groups. Brain of each fish was kept separate. Total RNA was extracted from the brain using TRIzol™ (Invitrogen, Carlsbad, CA, USA) method. At least four pituitary glands were pooled per group (n = 4 pools per treatment), and total RNA was extracted using a commercial kit (PureLink™ RNA Mini Kit, Ambion, Life Technologies, Carlsbad, CA, USA). After RNA extraction, the usual downstream procedures were followed according to methods described above. The relative mRNA levels of *gnrh2* and *gnrh3* (gonadotropin-releasing hormones), *gnih* (gonadotropin-inhibitory hormone), and *crf* (corticotropin-releasing hormone) were evaluated in the brain, and the *lhb* (luteinizing hormone), *fshb* (follicle-stimulating hormone), and *tsh* (thyroid-stimulating hormone) mRNA levels were determined in the pituitary gland. mRNA levels of the targets (Cts) were normalized by the reference gene *β-actin* and expressed as relative values of the control group (as fold induction) according to the 2 ^−(ΔΔCt)^ method. Primers were designed based on zebrafish sequences available at Genbank (NCBI) **(Table 1)**.

**Table 1.** Primers used for gene expression studies (qPCR). (FW = Forward; RV = Reverse).

### 2.6 Quantification of androgen (11-KT) plasma levels

Blood from adult male fish in different conditions (control, T4, methimazole, and methimazole + T4 groups) were collected (n = 8 per condition). Fish were euthanized, and the caudal peduncle was cut for blood sampling using heparinized syringes and tubes. Samples were centrifuged at 4°C for 10 min at 800 ×g (Eppendorf Centrifuge 5424 R), and 11-Ketotestosterone (11-KT) plasma levels were quantified by ELISA (582751, Cayman Chemical), following the manufacturer’s instructions. The results were evaluated as nanograms of 11-KT per milliliter of plasma.

### 2.7 Testis tissue culture

Testes were dissected and cultured using an *ex vivo* organ culture system described previously (Leal et al., 2009). For short-term incubations (18 h for 11-KT release analysis), testes were submerged in a culture medium, whereas for long-term exposure (7 days for histomorphometrical analysis and gene expression), testes were placed on a nitrocellulose membrane on top of a cylinder of agarosis and exposed to 1 mL of medium culture in 24-well flat-bottom plates, as described by Leal and collaborators (2009).

### 2.8 Short-term (18 h) incubation

Testes were collected from eight animals per condition (control, T4, methimazole, methimazole + T4) post-dissection. One testis was cultivated in the Lebovitz medium (L-15), whereas its contra-lateral one in L-15 containing recombinant zebrafish Fsh (rzfFsh; 100 ng/mL). Testes were incubated in 96-well plates containing 200 μL of solution in each well at 28 °C. Following incubation, testes were individually weighed, and the medium was collected and stored at −20 °C for androgen release (11-KT) assay (see sub-section 2.9).

### 2.9 *in vitro* 11-KT release by zebrafish testicular explants in short-term incubation

This technique was used to examine if treatment conditions (T4, methimazole, methimazole + T4) modulated rzfFsh (100 ng/mL)-induced androgen release by zebrafish testis. The androgen (11-Ketostesterone, 11-KT) release capacity of zebrafish testicular tissue into culture medium was measured after a short-term (18 h) *ex vivo* culture system as described previously (García-López et al., 2010). The levels of 11-KT released in the culture medium were quantified by ELISA using a commercial kit (Cayman Chemical) following manufacturer’s instructions.

### 2.10 Long-term (7 days) incubation

To study the effects of T3 (100 nM/mL) (n = 8) on zebrafish spermatogenesis, long-term incubations were performed. For that, each testis was placed on a nitrocellulose membrane measuring 0.25 cm^2^ (25 μm of thickness and 0.22 μm of porosity) on top of a cylinder of agarose (1.5% w/v, Ringer’s solution, pH 7.4) with 1 mL of culture medium into a 24-well plate. One testis was incubated in the presence of T3 alone and its contralateral one in a basal culture medium. The medium was changed every 3 days. After 7 days, testes were collected for histomorphometrical analysis. The proportion of section area occupied by different germ cell types were represented as fold-change of basal value. The relative number of spermatozoa per area was quantified as described above (see sub-section 2.4). Also, this technique was used to analyze if different concentrations of T3 and T4 (10, 100 and 1000 nM/mL) modulate expression of selected genes in zebrafish testis. For that testis were collected, total RNA was extracted from testis explants (n = 8) and the relative mRNA levels of *nanos2, sycp3l, 3β-HSD* (3-beta (β)-hydroxysteroid dehydrogenase), and *cyp17a1* were evaluated as described above (see sub-section 2.5) **(Table 1)**.

### 2.11 T3 injections

Adult zebrafish males were intracoelomically injected with 0 and 250 ng of T3 per fish (n = 16). The dose was selected based on their ability to induce deiodinase type 3 transcript as described previously (Nelson and Habibi, 2008; Nelson et al., 2010). First, a stock solution of T3 was prepared in 0.02 M sodium hydroxide and diluted in physiologic saline. The control group was injected with a vehicle (only physiologic saline solution). After twelve hours, the animals were euthanized, and brain and pituitary glands were collected. Brain of each fish (n = 8) was kept separate. The pituitaries were pooled in groups of four per group (n = 4 pools of 4 pituitaries per condition) for RNA extraction. mRNA release levels of *gnrh2* and *gnrh3*, *gnih*, and *crf* were measured in the brain, and *lhb*, *fshb*, and *tsh* were measured from pituitary glands **(Table 1)**. RNA extraction and downstream procedures were followed as previously described in sub-section 2.5.

### 2.12 Statistical analysis

All data were subjected to normality Shapiro-Wilk test, which was followed by the Bartlett homogeneity variance test. Significant differences were identified using unpaired or paired t-tests for two groups or one-way ANOVA followed by the Student–Newman–Keuls or Dunnett’s tests for three or more groups. Significance level (*p*) was considered at 0.05 in both cases. Data are presented as mean ± SEM (Standard Error of Mean). All data analysis was performed by Graph Pad Prism software 7.04 (*Graph Pad Software*, Inc., San Diego, CA, USA, http://www.graphpad.com).

## 3 Results

### 3.1 Thyroid follicles analysis

Thyroid gland follicles were examined histologically in the control group and following treatments with methimazole and methimazole + T4 **(Fig. 1(A)-(D))**. The control group thyroid follicles consisted of squamous/cuboidal epithelial cells filled with colloid **(Fig. 1(B))**. The results demonstrate a significant change in the histological condition of thyroid follicles in fish treated with methimazole. Three-week exposure to 1 mM methimazole resulted in follicles with columnar epithelium, follicle cell hypertrophy, and colloid depletion **(Fig. 1(C))**. These morphological changes were consistent with studies in which zebrafish were exposed to methimazole (Rodrigues et al., 2021), and other goitrogens, such as 6-n-propyl-2-thiouracil (PTU) (Van der Ven et al., 2006) and perchlorate (Patiño et al., 2003). The observed effect of methimazole was reversed by co-treatment with T4, in which the thyroid follicles were found to be morphologically similar to the control group **(Fig. 1(D)).** The results demonstrate that methimazole-induced hypothyroidism in zebrafish had altered thyroid function following treatment with the goitrogen. Also, the results demonstrate that co-treatment with T4 restored the zebrafish thyroid follicular structure.

**Figure 1:**
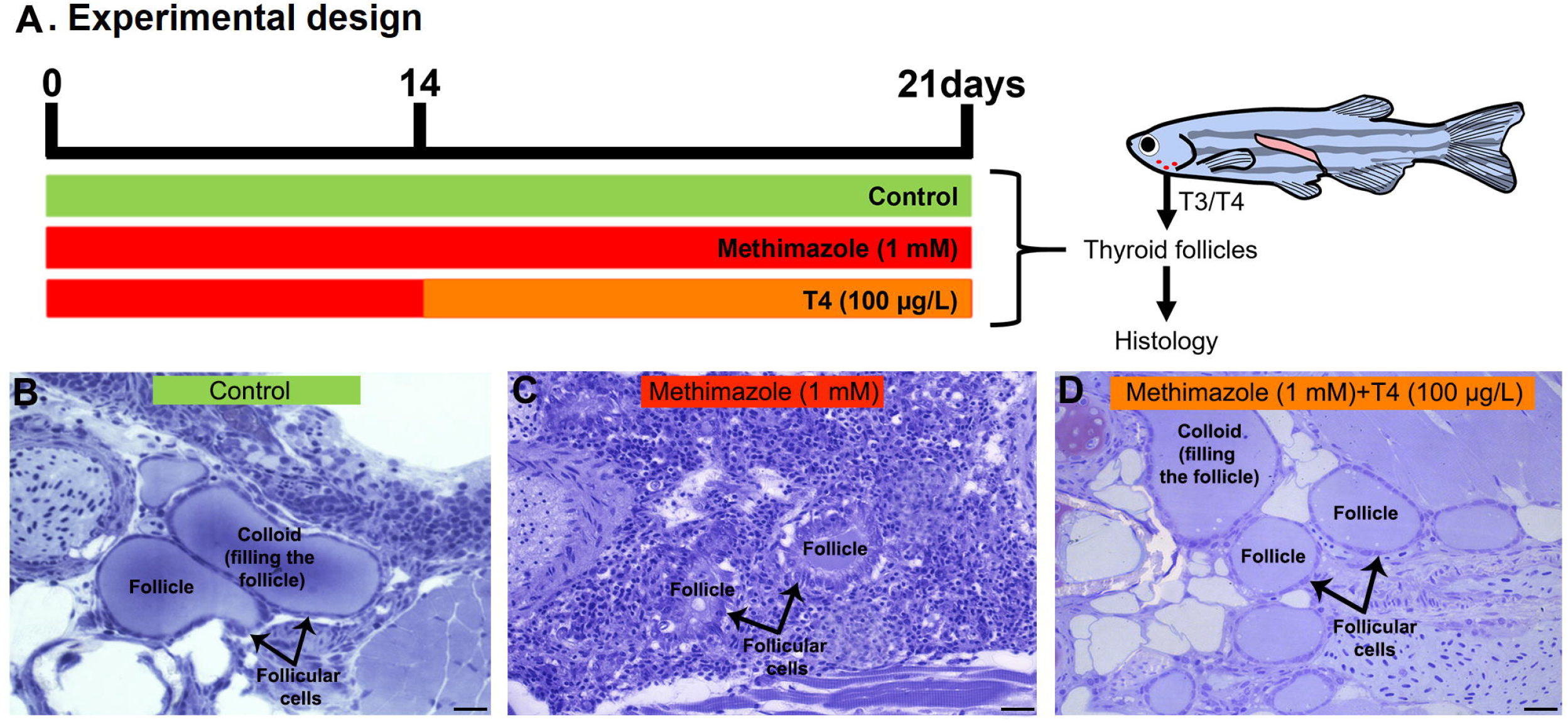
**(A)** Experimental design representation of treatments: control (non-treated fish), methimazole (1 mM) and methimazole (1 mM) + T4 (100 μg/L) groups. Zebrafish adult males were exposed to reconstituted water containing 1 mM methimazole (goitrogen) for three-weeks or 1 mM methimazole following T4 (100 μg/L) added in the water from the second week until the end of experiment. The control group received the same volume of vehicle solution. After 21 days heads were collected for histological analysis **(B-D)**. Control animals **(B)** have thyroid follicles with squamous/cuboidal epithelial cells and are completely filled with colloid. Fish treated with methimazole **(C)** showed follicles with columnar epithelium, follicle cell hypertrophy and colloid depletion, while fish co-treated with T4 **(D)** showed follicles similar to the control animals. Staining: Toluidine blue with sodium borate. Scale bar = 20 μm.

### 3.2 Methimazole-induced hypothyroidism and reversal treatment with T4: histomorphometrical analysis of the zebrafish testis

Methimazole-induced hypothyroidism promoted histomorphometrical changes in the proportion of germ cell cysts compared to the control **(Fig. 2(A)–(D))**. There was a significant increase in the proportion of the area occupied by type A undifferentiated spermatogonia (A_und*_), type A differentiated (A_diff_) and spermatogenic cysts containing type B spermatogonia (SpgB) in the methimazole group as compared to control **(Fig. 2 (D))**. The number of meiotic cells (Spc) and post-meiotic haploid cell population (Spt) did not change between control and methimazole-treated group **(Fig. 2 (D))**. However, the number of spermatozoa reduced drastically when compared to control as clearly evidenced in the photomicrographs and analysis of spermatozoa number by field (**Fig. 2(A), (B) and (E))**. Co-treatment with T4 rescued the proportion of A_und*_, A_diff_ and SpgB types returned to its basal values, while the proportion area occupied by Spc and Spt significantly increased **(Fig. 2 (D))**. Interestingly, the production of spermatozoa was recovered in the co-treatment with T4 (as viewed in the fields of **Fig. 2(A), (C) and (E)**).

**Figure 2:**
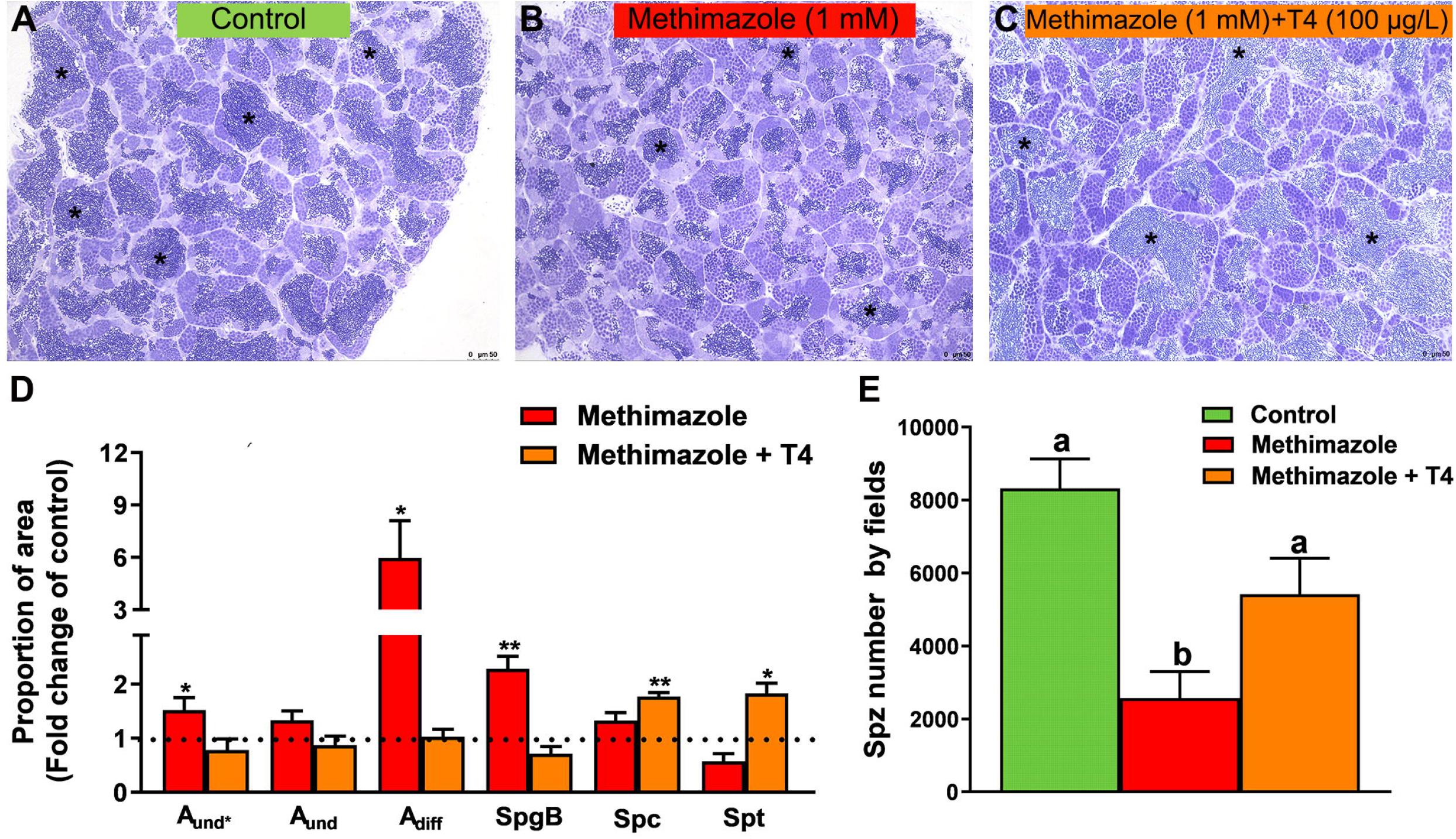
Histomorphometrical evaluation of zebrafish testes after *in vivo* exposure to methimazole and co-treatment with T4 for 21 days. Control group (non-treated fish) **(A)**. Methimazole-treated group **(B)**. Methimazole co-treated with T4 **(C)**. Asterisks in **(A)**, **(B)** and **(C)** indicate the testicular lumen with spermatozoa that appeared reduced in the methimazole group. **(D)** Proportion of section area occupied by spermatogenic cysts containing types A undifferentiated spermatogonia (A_und*_, A_und_), A differentiated spermatogonia (A_diff_), B spermatogonia (SpgB), spermatocytes (Spc), and spermatids (Spt). Bars (mean ± SEM; n = 8) are expressed as fold-change relative to the untreated group (control) (dotted line set at 1). **(E)** Spermatozoa number per field generated by using IMAGEJ Software from control and treatments. ANOVA followed by Dunnett’s multiple comparison tests. Different letters denote significant differences (*p* < 0.05) between different treatment conditions with the control. Asterisks denote statistical significance differences between control, methimazole and methimazole + T4 groups; * *p < 0.05;* ** *p < 0.01* (Student unpaired t-test; n = 8). Staining: Toluidine blue. Scale bar = 50 μm.

### 3.3 Methimazole and co-treatment with T4: testicular transcript levels

Transcript levels of several genes involved in reproduction were measured by qPCR in the testis from methimazole-induced hypothyroidism and rescued groups (methimazole + T4) **(Fig. 3)**. In this study we measured transcript levels of two thyroid hormone receptor subtypes (*thrα, thrβ*). The *thrα* transcript level was higher in the methimazole treated group than control, but the difference was not statistically significant **(Fig. 3(A)).** The *thrα* transcript level was further increased significantly in the methimazole + T4 treated group **(Fig. 3(A)).** Similarly, the *thrβ* was increased in the methimazole and methimazole + T4 treated groups, compared to the control **(Fig. 3(B))**.

**Figure 3:**
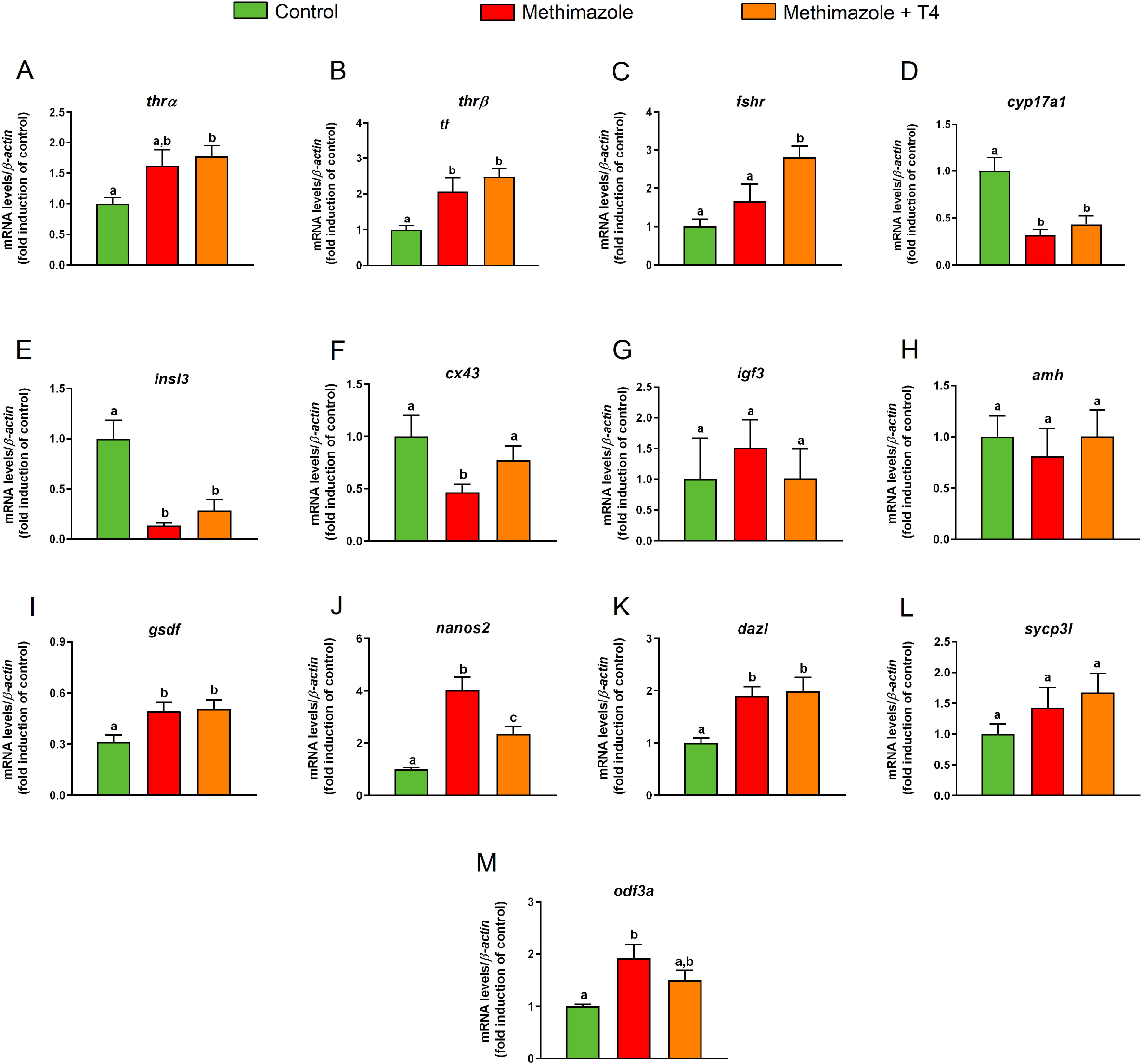
Relative mRNA levels of several genes expressed in zebrafish testis after *in vivo* exposure to methimazole and methimazole co-treated with T4 for 21 days. The selected genes *thrα* and *thrβ* (thyroid hormones receptor) **(A-B)**; *fshr* (follicle-stimulating hormone receptor) **(C)**; genes expressed by somatic cells (Leydig and Sertoli cells) **(D–I)**; *cyp17a1* (17α-hydroxylase/17,20 lyase) **(D)**; *insl3* (insulin-like peptide 3) **(E)**; *cx43* (testicular connexin) **(F)**; *igf3* (insulin-like growth factor 3) **(G)**; *amh* (anti-Müllerian hormone) **(H)**; *gsdf* (gonadal somatic cell derived factor) **(I)**; and germ cell markers **(J–M)**; *nanos2* **(J)**; *dazl* (deleted-in azoospermia-like) **(K)**; *sycp3l* (synaptonemal complex protein 3) **(L)**; *odf3a* (outer dense fiber of sperm tails 3B) **(M)** were evaluated. Ct values were normalized with *β-actin* and expressed as relative values of control levels of expression. Bars represent the mean ± SEM fold change (n = 8) relative to the control, which is set at 1. Student unpaired t-test. Different letters denote significant differences (*p* < 0.05) between different treatment conditions with the control.

We also measured mRNA levels of *fshr* which was not affected in the methimazole treated group, but significantly increased in the methimazole + T4 treated fish, compared to the control **(Fig. 3(C))**.

In the present study we measured transcript levels for *cyp17a1* and insulin-like peptide 3 (*insl3*) genes. Treatment with methimazole significantly reduced the *cyp17a1* and *insl3* transcript levels compared to control **(Fig. 3(D) and (E))**. Co-treatment with T4 did not influence the methimazole induced response on the expression of these transcripts **(Fig. 3(D) and (E))**. We also observed significant reduction in the testicular connexin (*cx43*) mRNA levels in the methimazole treated group, compared to control **(Fig. 3(F))**. Co-treatment with T4 in this case increased the *cx43* mRNA to a level not significantly different from the control **(Fig. 3(F))**.

Among others, *igf3, amh*, and *gsdf* genes are known to be expressed in the Sertoli cells. Treatment with methimazole or methimazole + T4 were without significant effects on *igf3* and *amh* transcript levels **(Fig. 3(G) and (H))**. The *gsdf* mRNA level, however, was higher in the methimazole and methimazole + T4 treated groups **(Fig. 3(I))**.

With regard to germ cell marker genes, such as marker for type A undifferentiated spermatogonia, *nanos2* mRNA level was significantly higher in the methimazole treated group compared to control **(Fig. 3(J))**. Co-treatment with T4, significantly reduced the methimazole-induced response to a level higher than the basal control **(Fig.3(J))**. The *dazl* (deleted-in azoospermia-like) transcript level expressed by A_diff_ and SpgB, increased in the methimazole treated group compared to the control **(Fig. 3(K))**. Co-treatment with T4 did not influence the methimazole-induced response **(Fig. 3(K))**. In our study, synaptonemal complex protein 3 (*sycp3l*) which is a marker for spermatocytes was not significantly affected by methimazole or methimazole + T4 treatments **(Fig. 3(L))**. However, the transcript level of the outer dense fiber protein 3 (*odf3a*) which is a marker for spermatids was increased following treatment with methimazole **(Fig. 3(M))**. Co-treatment with T4 reduced the methimazole-induced response to a level not significantly different from the basal control **(Fig. 3(M))**.

### 3.4 Measurement of 11-KT levels

In the present study, we measured the 11-KT concentration to partially assess the effect of methimazole-induced hypothyroidism on steroidogenesis, *in vivo* and *in vitro* **(Fig. 4(A))**. Four treatment groups were studied in this experiment, including control, control + T4, methimazole and co-treatment (methimazole + T4) groups **(Fig. 4(A))**. Animal exposure with T4 (control+T4) significantly reduced the plasma 11-KT level as compared to control levels **(Fig. 4(B))**. Also, treatment with methimazole significantly reduced the plasma 11-KT concentration to almost undetectable level compared to the control level of over 40 ng/mL **(Fig. 4(B))**. However, co-treatment with T4 significantly increased and nullified the methimazole-induced response to a level not significantly different from the control **(Fig. 4(B))**.

**Figure 4:**
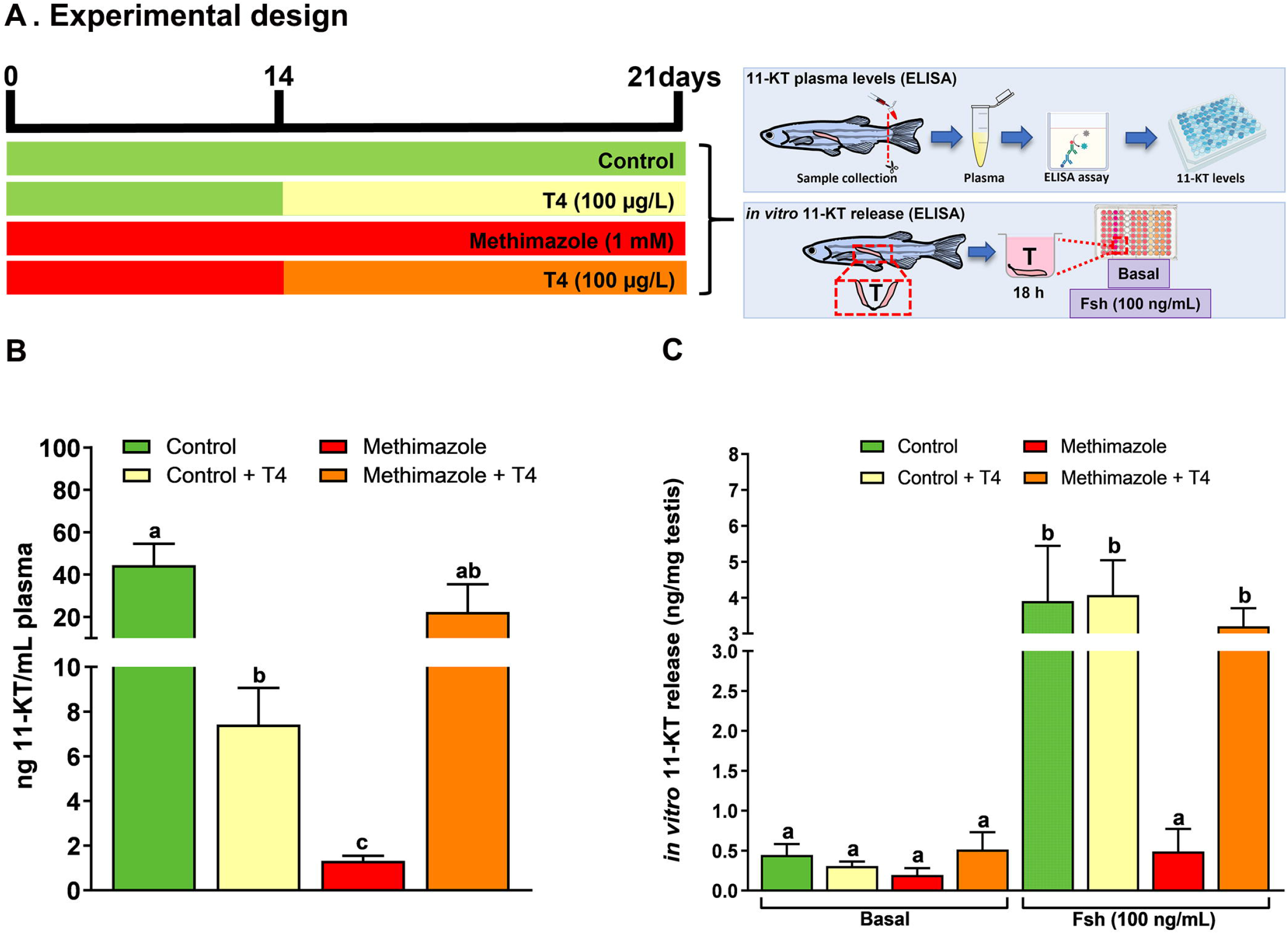
**(A)** Experimental design. 11-Ketotestosterone (11-KT) plasma levels (ng/mL plasma) was measured in zebrafish males exposed to different treatments: Control (non-treated fish), T4 (100 μg/L), methimazole (1 mM) and methimazole (1 mM) + T4 (100 μg/L). Amounts of 11-KT (ng/mg of testis weight) released by zebrafish testes were measured in the incubation media after short-term exposure (18 h) in the presence or absence of Fsh (100 ng/mL) from control, T4, methimazole and methimazole + T4 groups. **(B)** Effect of T4, methimazole and combination of methimazole and T4 on 11-KT plasma levels. **(C)** Androgen (11-KT) release from zebrafish testicular explants previously treated with T4, methimazole or methimazole co-treated with T4. Bars represent the mean ± SEM (n = 8). ANOVA followed by Tukey’s test. Different letters denote significant difference (*p* < 0.05) between different treatments compared to the respective control group.

We also measured the androgen (11-KT) release capacity of zebrafish testicular tissue using the *ex vivo* culture system **(Fig. 4(A))**. In this experiment, we compared testis taken from control and those exposed to T4 (100 μg/L), methimazole (1 mM) and methimazole co-treated with T4. We tested the 11-KT release response into culture media following *in vitro* treatment for 18 h to recombinant zebrafish Fsh (100 ng/ml) **(Fig. 4(A))**. In the basal medium condition, the treatments (T4, methimazole and methimazole + T4) did not change the basal androgen (11-KT) release capacity of zebrafish testicular tissue **(Fig. 4(C))**. However, as expected, treatment with Fsh significantly increased the 11-KT level in the control group **(Fig. 4(C))**. Also, treatment with T4 was responsive to Fsh but to a level not significantly different from the control **(Fig. 4(C))**. In comparison, the isolated testis from methimazole-treated zebrafish was completely unresponsive to Fsh **(Fig. 4(C))**. However, co-treatment with T4 restored the Fsh-induced response by increasing the 11-KT concentration to the level observed following treatment of the control group with Fsh alone **(Fig. 4(C))**.

### 3.5 Transcript levels of selected genes in the brain and pituitary

In this experiment, we measured brain transcript levels of a number of neurohypothalamic peptides, including *gnrh2* and *gnrh3*, *gnih*, and *crf*, as well as pituitary gonadotropin hormone subunits, *fshb, lhb*, and *tshb* in the methimazole-induced hypothyroidism in fish **(Fig. 5(A) and (C))**. As shown in **Fig. 5(C)**, we observed a significant increase in the *gnih* transcript level compared to the control shown as the dotted line. Transcript levels of the other neuropeptides measured including *gnrh2*, *gnrh3* and *crf* remained unchanged. In the same group of methimazole-treated fish, the results the pituitary glycoprotein hormone subunits, demonstrate significant increase in *fshb* mRNA and a massive increase in *tshb* transcript levels in the methimazole-induced hypothyroid fish **(Fig. 5(C))**. No change was observed in the *lhb* transcript level **(Fig. 5(C))**.

**Figure 5:**
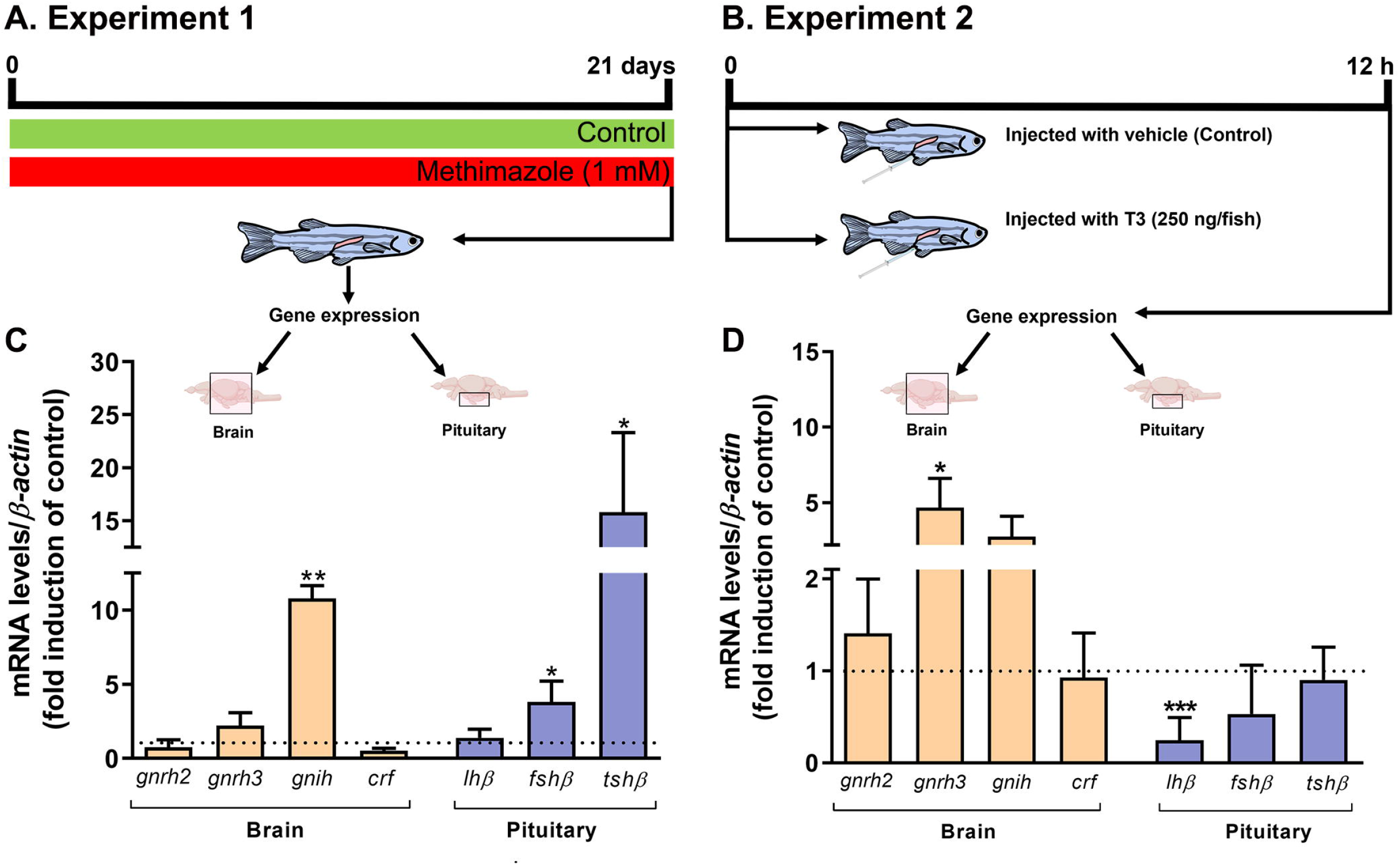
**(A)** Experiment 1: zebrafish males were exposed to methimazole-induced hypothyroidism for 21 days. **(B)** Experiment 2: zebrafish males were injected with 0 or 250 ng of T3/fish and tissue were collected 12 h post-injection. Relative mRNA levels of selected genes, including *gnrh2* and *gnrh3* (gonadotropin-releasing hormones), *gnih* (gonadotropin-inhibitory hormone), and *crf* (corticotropin-releasing hormone), expressed in the brain (n = 8), and *lhβ* (luteinizing hormone), *fshβ* (follicle-stimulating hormone), and *tshβ* (thyroid-stimulating hormone) expressed in the pituitary (n = 4 pools of 4 pituitaries for each pool) from the methimazole group **(C)** and 12 h post-injection with 0 and 250 ng of T3/fish **(D)**. Ct values were normalized with *β-actin* and expressed as relative values of control (0 ng/mL) levels of expression. Bars represent mean ± SEM fold-change relative to the control, which is set at 1. Student unpaired t-test, *\*p* < 0.05, *\*\*p < 0.01* and *\*\*\*p* < 0.001.

We also measured the same transcript levels following 12 hours acute treatment with 250 ng of T3/fish *in vivo* **(Fig. (B) and (D))**. In this experiment T3 injection did not significantly alter *gnih, gnrh2* and *crf*, but significantly increased the *gnrh3* transcript level **(Fig. 5(D))**. In the same group of fish, acute treatment with 250 ng of T3/fish significantly reduced the pituitary *lhb* but was without effect on the *fshb* and *tshb* transcript levels **(Fig. 5(D))**.

### 3.6 *in vitro:* effects of T3 on zebrafish spermatogenesis

In this study, we also investigated the direct action of T3 at 100 nM/mL on *ex vivo* culture of zebrafish testis for 7 days **(Fig. 6(A))** and results provide information on the proportion of germ cells in the cultured testis. Histomorphometrical evaluation of zebrafish testis revealed that treatment with T3 significantly stimulated the type A undifferentiated spermatogonia (A_und_) abundance with no effect on type A differentiated spermatogonia cells as compared to basal medium incubation **(Fig. 6(B))**. In the same tissue, we observed a reduction in type B spermatogonia (SpgB) abundance **(Fig. 6(B))**. As for meiotic and post-meiotic cells, treatment with T3 reduced the proportion of area occupied by Spc and Spt when compared to basal condition **(Fig. 6(B))**. Interestingly, the number of spermatozoa was stimulated in zebrafish testis treated with T3 as clearly shown by the analysis of spermatozoa number by field **(Fig. 6(C))**. In addition to these data, we evaluated the effects of thyroid hormones on gene expression **(Fig. 6 (D)-(G)) (see Supplementary Figure)**. Different concentrations of T3 and T4 (10, 100 and 1000 nM) were tested in zebrafish testis. T4 did not modulate the expression of any transcript (*nanos2, sycp3l, 3β-HSD* and *cyp17a1*) **(see Supplementary Figure)**. On the other hand, expression analysis showed that T3 (100 and 1000 nM/mL) increased *nanos2* **(Fig. 6 (D))**. Interestingly, T3 also stimulates the *sycp3l*, the meiotic marker expression (100 nM/mL) **(Fig. 6 (E))**. Levels of *3β-HSD* and *cyp17a1* were also measured and did not change within any levels of T3 **(Fig. 6(F)-(G))**.

**Figure 6:**
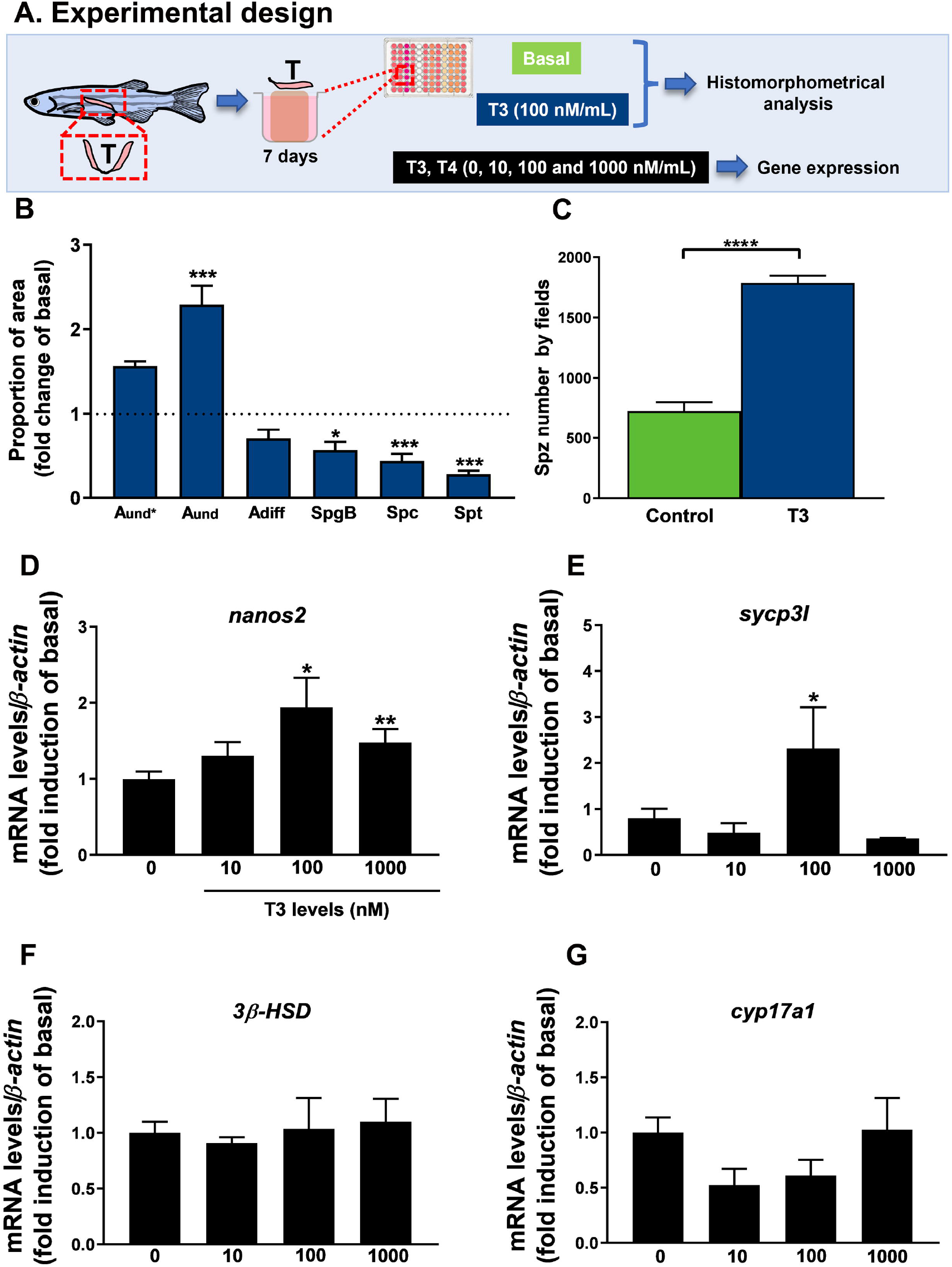
**(A)** Experimental design of histomorphometrical analysis of zebrafish testicular explants incubated for 7 days (long-term exposure) with T3 (100 nM/mL) compared to the control (basal) and gene expression of relative mRNA levels of several selected genes in zebrafish testis incubated for 7 days to different concentrations of T3 (0, 10, 100 e 1000 nM/mL). **(B)** Histomorphometrical evaluation of testicular explants containing types A undifferentiated spermatogonia (A_und*_, A_und_), type A differentiated spermatogonia (A_diff_), type B spermatogonia (SpgB), spermatocytes (Spc), and spermatids (Spt). Bars (mean ± SEM; n = 8) are expressed as fold-change relative to the untreated group (control) (dotted line set at 1). **(C)** Spermatozoa number per field generated by using IMAGEJ Software from zebrafish explants incubated for 7 days with basal (L-15) and T3 (100 nM/mL). Bars (mean ± SEM; n = 8). Student paired t-test, * *p* < 0.05 and \*\*\**p* < 0.001 denote significant differences between control and treated fish. The selected genes *nanos2* **(D)**, *sycp3l* (synaptonemal complex protein 3) **(E)**, *3β-HSD* (3-beta (β)-hydroxysteroid dehydrogenase) **(F)**, and *cyp17a1* (17α-hydroxylase/17,20 lyase/17,20 desmolase) **(G)** were evaluated. Ct values were normalized with *β-actin* and expressed as relative values of basal (0 ng/mL) levels of expression. Bars represent the mean ± SEM fold change (n = 8), relative to the control (basal, 0 ng/mL), which is set at 1. Paired *t - test, *p* < 0.05, *\*\*p < 0.01*.

## 4 Discussion

This study demonstrated the importance of thyroid hormones as a factor controlling zebrafish spermatogenesis, using methimazole-induced hypothyroidism in fish as a model organism. Several other studies have used antithyroid agents to investigate the effects of thyroid hormones on vertebrate reproduction, including fish (Swapna et al., 2006; Weng et al., 2007; Sharma and Patiño, 2013; Sharma et al., 2016; Tovo-Neto et al, 2018; Figueiredo et al., 2019; Rodrigues et al., 2021). In our study, the efficacy of hypothyroidism was considered based on histological conditions of thyroid follicles. Our results demonstrated that thyroid follicles clearly disrupted following a three-week exposure to 1 mM methimazole. However, the normal thyroid follicles were recovered after co-treatment with T4. Also, testicular connexin (*cx43*) levels as it is the potential target for thyroid hormones (Poguet et al., 2003; Gilleron et al., 2006). Interestingly, Gilleron and collaborators (2006) demonstrated that propylthiouracil (PTU; another thyroid disruptor) decreases *cx43* levels in rats. In our study, the lower expression of *cx43* in the methimazole group indicates that methimazole treatment led to hypothyroidism in the male zebrafish.

The present study provides clear evidence that hypothyroidism disrupts germ cell development in the zebrafish testis. Plasma concentration of androgen (11-KT) levels was significantly lower in the hypothyroid group when compared to that in the euthyroid group. Moreover, the relative mRNA levels of selected testicular genes expressed by somatic and germ cells were altered in hypothyroidism, and in addition to decreased 11-KT plasma levels, hypothyroidism induced by methimazole also impaired genes expressed in the brain and pituitary gland. We used co-treatment with T4, which is the primary secretory product of the thyroid gland, to potentially reverse the effects of methimazole treatment. In addition to these *in vivo* experiments, we also examined the acute effects of thyroid hormones in zebrafish testicular explants in the presence or absence of rzfFsh in different treatments. Thus, this study provides a novel and practical approach to better understand the nature of thyroid hormone actions on the HPG axis in male zebrafish.

Histomorphometric evaluation of zebrafish spermatogenesis demonstrated that methimazole-induced hypothyroidism remarkably increased the proportion area occupied by type A_und*_ (the most undifferentiated spermatogonia), type A_diff_ and type B spermatogonia, while the number of spermatozoa was decreased. Furthermore, our results demonstrate that the proportion of Spc and Spt were not affected in hypothyroid testis compared with the control animals. It has been well-established that spermatogenesis development is supported and regulated by gonadotropic hormones, FSH and LH which mainly target actions on somatic testicular cells such as Sertoli and Leydig cells (Planas et al., 1993; Campbell et al., 2003; Huhtaniemi and Themmen, 2005; García-López et al., 2010; Schulz et al., 2010; Flood et al., 2013). Likewise, thyroid hormones have also been shown to play an essential role in testicular physiology (Chiao et al., 2002; Cooke et al., 2004; Holsberger and Cooke, 2005; Wagner et al., 2008; 2009; Morais et al., 2013; Tovo-Neto et al., 2018). In rats, previous studies have reported that hypothyroidism decreases plasma levels of gonadotropins and reduces the number and size of gonadotrophs (Bruni et al., 1975; Ruiz et al., 1988), suggesting that the absence of thyroid hormones is associated with gonadal dysfunctions (Amin and El-Sheikh, 1977; Hernandez, 2018). Further, the level of relevant androgen (11-KT) that stimulates spermatogenesis decreased significantly in the plasma of zebrafish following treatment with methimazole. These results, in part, could explain the observed increase in different populations of spermatogonia (A_und*_, A_diff_ and B) and decrease in spermatozoa number, demonstrating the impact of thyroid hormone depletion on germ cell development and testicular function. Similar findings have been reported for male Japanese quail, wherein treatment with methimazole decreased body and testes weight as well as plasma levels of LH and testosterone (Weng et al., 2007). In this same study, the data showed decreased spermatogenesis in seminiferous tubules of the treatment group (Weng et al., 2007). Altogether, these results demonstrate that gonadal function is associated with normal thyroid action. Fluctuations in thyroid hormone levels can directly modulate gonadotropin actions and affect Sertoli and Leydig cell proliferation (Castañeda-Cortés et al., 2014), resulting in the impairment of spermatogenesis, which may be caused by low FSH level and delay of Sertoli cell maturation (Hernandez, 2018).

We also examined the effects of methimazole co-treated with T4. Our results demonstrate that co-treatment with T4 partially restored zebrafish spermatogenesis and spermatozoa production in the methimazole-induced hypothyroid fish. Interestingly, the administration of T4 to the hypothyroid group increased the proportion of meiotic (Spc) and post-meiotic (Spt) cells compared with the control and methimazole groups. These results support the hypothesis that thyroid hormones are involved, directly or indirectly, with meiosis entry in the zebrafish testis.

The observed transcript analysis in zebrafish testis is to some extent in agreement with histomorphometric results. With respect to the germ cell markers genes of zebrafish spermatogenesis, the results indicated that mRNA levels of *nanos2* (marker of type A undifferentiated spermatogonia) (Aoki et al., 2009) and *dazl* (marker of differentiation) (Chen et al., 2013) were higher in the methimazole group, in agreement with the histomorphometry showing an accumulation of pre-meiotic cells (A_und*_, A_diff_ and B). The observed upregulation of these transcripts in the methimazole treated group suggest that euthyroid condition may be important for normal spermatogonial cell development in zebrafish. However, the level of *odf3a* (marker of spermatids) (Yano et al., 2008) was upregulated in the methimazole group, although the proportion of spermatids was not affected in the present study. The affected testicular functions did not always recover following co-treatment with T4 to counter hypothyroidism.

Other important components of the thyroid hormones axis are the thyroid receptors (TRs), which are crucial for testis development and function (Valadares et al., 2008; Dittrich et al., 2011; Lema et al., 2009; Wajner et al., 2009). According to Habibi et al., (2012), TRs are expressed by several different cell types, and thyroid hormones have pleiotropic effects, including effects on the gonads. In the latter study in goldfish, *in vivo* and *in vitro*, thyroid hormones were shown to exert both direct and indirect actions on gonadal steroid synthesis, and steroid receptor expression in a seasonally dependent manner (Allan and Habibi, 2012; Habibi et al., 2012). In zebrafish testis, *thrα* was shown to be expressed in the Sertoli and Leydig cells, whereas *thrβ* expression was only observed in the Leydig cell (Morais et al., 2013). In the present study, methimazole-induced hypothyroidism stimulated *thrβ* transcript level, but increase in *thrα* was only significant in presence of T4 compared to control. Other investigators also reported that T3 treatment only elevated *thrβ* in the ovary and testis of *Pimephales promelas* (Lema et al., 2009). Similarly, *fshr* mRNA levels were also upregulated in hypothyroid co-treated with T4. These results suggest that chronic hyperthyroidism may alter thyroid hormone sensitivity and possibly alter other parameters not clear at the present time. Similarly, in rats, higher FSH-R mRNA levels were detected in hypothyroidism. It was suggested that the upregulated FSH-R mRNA level may have resulted from the elevated proportion of Sertoli cells in rats (Rao et al., 2003). Likewise in zebrafish, the increase of *fshr* transcripts following co-treatment with T4 could be due to Sertoli cell proliferation. Morais et al., (2013) showed that T3 stimulates *in vitro* Sertoli cell proliferation in zebrafish testes. Moreover, in zebrafish testis, Sertoli cells express both *thrα* and *fshr*, and treatment with T3 potentiates Fsh action in the zebrafish testis *in vitro* (Morais et al., 2013). This is also consistent with the observation that *fshr* transcript level was stimulated by methimazole-induced hypothyroid fish co-treated with T4. These results support the view that thyroid hormones and gonadotropins stimulate spermatogenesis by stimulating gonadal androgen production, which in turn mediate the start of spermatogenesis.

Interestingly, mRNA expression of other key gonadal growth factors as *igf3* (Wang et al., 2008) and *amh* (Miura et al., 2002) did not change in the methimazole or methimazole co-treated with T4 groups. However, *gsdf* (Gautier et al., 2011) was upregulated in both groups. *gsdf* is expressed by Sertoli cells and exerts a crucial role in the control of germ cell proliferation and differentiation in spermatogenesis (Gautier et al., 2011; Yan et al., 2017).

Decreased thyroid hormones levels by methimazole in male zebrafish are associated with decreased 11-KT plasma levels. Previous studies have shown that PTU treatment also decreases serum testosterone concentrations in rats (Chiao et al., 2000). In another study, exposure of *Clarias gariepinus* to thiourea, reduced androgen levels leading to testicular regression (Swapna et al., 2006; Swapna and Senthikumaran, 2007). This is consistent with the present study demonstrating lower 11-KT plasma levels in the methimazole group, in addition to lower levels of steroidogenic enzyme (*cyp17a1*) and androgen-sensitive gene (*insl3*) mRNA levels. These transcript levels remained the same even after co-treatment with T4. However, treatment of methimazole-induced hypothyroid fish effectively rescued 11-KT levels in the plasma which was correlated with the revival of spermatozoa number in zebrafish testis. Thus, impairing thyroid action within the physiological limit of restoration is a valid approach to explain the role of thyroid hormones in male zebrafish reproduction. Interestingly, Houbrechts and collaborators (2019) demonstrated the Dio2-knockout (KO) zebrafish (*dio2^-/-^*) strongly decreased androgen (11-KT and testosterone) levels in the testis, and steroidogenesis and steroid signaling were similarly affected when compared with control animals. This indicates that absence of thyroid hormones by goitrogen treatment or DIO2 KO, which suppresses thyroid hormone production, may act directly on the testis to repress steroidogenesis. These data also indicate that normal thyroid hormone levels are crucial to normal reproduction in zebrafish.

The result in this study helps to elucidate the mechanisms underlying the observed reduction in androgen level in male zebrafish following treatment with methimazole. While transcript levels of *gnrh2* and *gnrh3* did not change, *gnih* mRNA levels were significantly elevated in the brain of animals after methimazole exposure. In fish, GnIH orthologs have double mechanism, including stimulatory and inhibitory actions on gonadotropin secretion and growth hormone production, depending on the species studied, mode of treatment, and season (Amano et al., 2006; Zhang et al., 2010; Moussavi et al., 2012; 2013; 2014; Branco et al., 2018; Fallah et al., 2019; Ma et al., 2020a, b). In addition, Tsutsui and collaborators (2017) showed that hypothyroidism induced by PTU increases GnIH expression and reduces gonadotropins and plasma steroid levels in female mice. The present results support the hypothesis that decrease in plasma androgen level in methimazole group is associated with the inhibition of the hypothalamic–pituitary-gonadal axis mediated, in part, by increased GnIH. As shown previously in goldfish, GnIH mediated inhibition of reproduction does not always correlate closely by inhibition of gonadotropin transcript levels due to uncoupling of release and synthesis. In this context, it was demonstrated that GnIH can increase gonadotropin subunit mRNA levels while reducing secretion of the hormones (Moussavi et al., 2012; 2013; 2014; Ma et al., 2020a, b). Thus, the inhibition may be at the protein level, since *fshb* transcript level was increased in the pituitary of the methimazole-treated group. Similarly, Yoshiura and collaborators (1999) found in goldfish that thiourea treatment per two weeks increased Fsh mRNA levels. A contributing factor could be due to feedback exerted by lower plasma 11-KT levels seen in these animals (Rojdmark et al., 1988; Trudeau et al., 1993; Habibi and Huggard-Nelson, 1998; Huggard-Nelson et al., 2002). There is increasing evidence that the effects of thyroid hormones as well as brain-pituitary-gonadal hormones are diverse, and change with season, mode of action, time-course, and concentration (Nelson and Habibi 2006, 2008, 2009; Nelson et al., 2011; Habibi et al., 2012; Moussavi et al., 2012, 2013, 2014; Nelson and Habibi 2016; Ma et al., 2020a, b). This is reflected in the present study as acute treatment for 12 hours with T3 injection resulting in a different effect than chronic treatments with T4 and methimazole on the hypothalamic–pituitary-gonadal axis. Injection with T3 increased *gnrh3* mRNA levels and either decreased (lhb) or had no effects (*fshb, tsh*) on the pituitary glycoprotein hormone subunit transcript levels. This is consistent with previous study in goldfish demonstrating reduction in gonadotropins and synthesis of gonadal steroids following 12 hour injection with T3 (Nelson et al., 2010).

In addition to these data, we evaluated the effects of thyroid hormones on zebrafish spermatogenesis. Histomorphometrical analysis showed an increase of type A undifferentiated spermatogonia (A_und_) proportion in T3 *in vitro* exposure (100 nM/mL), which may reflect the up-regulation of *nanos2* (marker of type A undifferentiated spermatogonia) (100 and 1000 nM/mL of T3). The number of spermatozoa also increased. This data corroborates the histomorphometry of the zebrafish testes from *in vivo* exposures using methimazole and methimazole co-treated with T4, indicating, therefore, a direct action of thyroid hormones in the zebrafish spermatogenesis. Previously, Morais et al. (2013) reported that the mitotic indices of A_und_ and Sertoli cells were stimulated after T3 treatment. Also, Safian et al. (2016) demonstrated that T3 enhanced the formation of new cysts of A_und_, accumulation of differentiated spermatogonia (A_diff_) and reduction of spermatogonia B. Morais et al. (2013) using lower dose (50 ng/mL) reported an increase of type A undifferentiated spermatogonia, which was corroborated by higher BrdU mitotic index of this cell type in T3-treated tissue. In this study, we also examined the effects of T3 and T4 on gene expression. Different concentrations of T3 and T4 (10, 100 and 1000 nM) were tested in zebrafish testis. Interestingly, T3 also stimulated the *sycp3l*, the meiotic marker expression (100 nM/mL). Levels of *3β-HSD* and *cyp17a1* were also measured and did not change within any level of T3. Another interesting result is that T4 does not exert any change. Thyroid hormones act via T3 in zebrafish testis. Altogether this data indicates that thyroid hormones (T3) stimulate zebrafish spermatogenesis by proliferation, differentiation, and meiosis. These results supply important data about the effects of thyroid hormones on brain-pituitary-testis and support interaction between reproductive and thyroid axes via neurohypothalamic peptides and gonadotropins (e.g. Fsh), which needs T3 to promote its functions in zebrafish testis.

## 5 Conclusion

The present study clearly demonstrated that hypothyroidism induced by methimazole significantly impair the progression of zebrafish spermatogenesis by inducing the accumulation of pre-meiotic cells, delaying differentiation and meiosis, and reducing the number of spermatozoa as well as impairing the hypothalamic–pituitary axis. We also provide evidence that testicular function is dependent on thyroid hormones. Taken together, these results provide support for that hypothesis that thyroid hormones are essential for spermatogenesis and maintaining normal function of the hypothalamic–pituitary-gonadal axis in adult zebrafish.

## Author Contributions

R.H.N., M.S.R., A.T.N., and H.R.H. designed the study; M.S.R., A.T.N., I.F.R., L.B.D., and H.P.F performed the experiments; All authors analyzed the data; R.H.N., M.S.R., A.T.N., and H.R.H. wrote the paper. All authors edited the article.

---

**Supplementary Figure:**
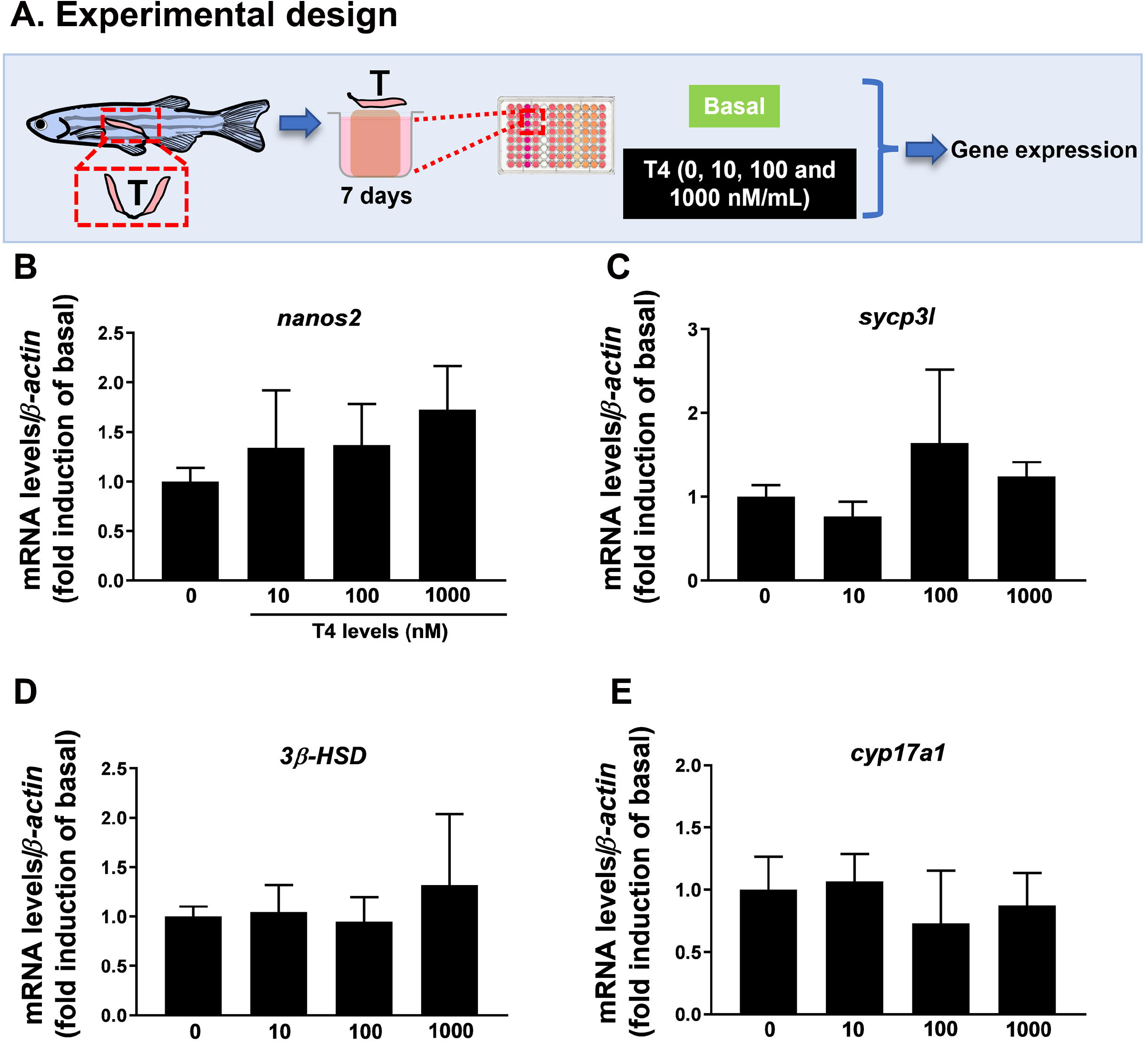
**(A)** Relative mRNA levels of several genes expressed in zebrafish testis after incubated for 7 days to different concentrations of T4 (0, 10, 100 e 1000 nM/mL). The selected genes *nanos2* **(B)**, *sycp3l* (synaptonemal complex protein 3) **(C)**, *3β-HSD* (3-beta (β)-hydroxysteroid dehydrogenase) **(D)**, and *cyp17a1* (17α-hydroxylase/17,20 lyase/17,20 desmolase) **(E)** were evaluated. Ct values were normalized with *β-actin* and expressed as relative values of basal (0 ng/mL) levels of expression. Bars represent the mean ± SEM fold change (n = 8), relative to the control (basal, 0 ng/mL), which is set at 1. Paired *t-test*.

